# Toward Machine Learning-based Data-driven Functional Protein Studies: Understanding Colour Tuning Rules and Predicting the Absorption Wavelengths of Microbial Rhodopsins

**DOI:** 10.1101/226118

**Authors:** Masayuki Karasuyama, Keiichi Inoue, Hideki Kandori, Ichiro Takeuchi

**Affiliations:** Nagoya Institute of Technology / JST PRESTO / National Institute for Materials Science; Nagoya Institute of Technology / OptoBioTechnology Research Center / JST PRESTO; Nagoya Institute of Technology / OptoBioTechnology Research Center; Nagoya Institute of Technology / RIKEN / National Institute for Materials Science

**Author notes:** Equally contributed (Masayuki Karasuyama, Keiichi Inoue).

## Abstract

The light-dependent ion-transport function of microbial rhodopsin has been widely used in optogenetics for optical control of neural activity. In order to increase the variety of rhodopsin proteins having a wide range of absorption wavelengths, the light absorption properties of various wild-type rhodopsins and their artificially mutated variants were investigated in the literature. Here, we demonstrate that a machine-learning-based (ML-based) data-driven approach is useful for understanding and predicting the light-absorption properties of microbial rhodopsin proteins. We constructed a database of 796 proteins consisting of microbial rhodopsin wildtypes and their variants. We then proposed an ML method that produces a statistical model describing the relationship between amino-acid sequences and absorption wavelengths and demonstrated that the fitted statistical model is useful for understanding colour tuning rules and predicting absorption wavelengths. By applying the ML method to the database, two residues that were not considered in previous studies are newly identified to be important to colour shift.

## 1 Introduction

Microbial rhodopsin is a photoreceptive membrane protein of microbial species, such as eubacteria, archaea, fungi, and algae. The functions of microbial rhodopsin are very diverse. Light-driven ion (proton, chloride, sodium, and so on) pumps, light-gated cation and anion channels, photochromatic gene regulator and light-regulated enzymes have been reported for various species^1^. The light-dependent ion-transport function of microbial rhodopsin is widely used in optogenetics for optical control of neural activity in the brain network^2^. Most microbial rhodopsins bind a common chromophore, all-*trans* retinal, via a protonated Schiff-base linkage in the center of the hepta-transmembrane scaffold (Fig. 1). Each microbial rhodopsin exhibits a variety of specific visible absorption wavelengths of their retinal. While the protonated all-*trans* retinal Schiff-base shows the absorption peak at ~450 nm in organic solvents^3^, the wavelengths of absorption maxima of retinal (λ_max_s) in microbial rhodopsin range from 436 nm of channel-rhodopsin from *Tetraselmis striata* (*Ts*ChR)^4^ to 587 nm of sensory rhodopsin 1^5^. This wide-range colour tuning of the retinal in rhodopsin is considered to be achieved by optimizing the steric and/or electrostatic interaction with surrounding amino-acid residues.

**Figure 1:**
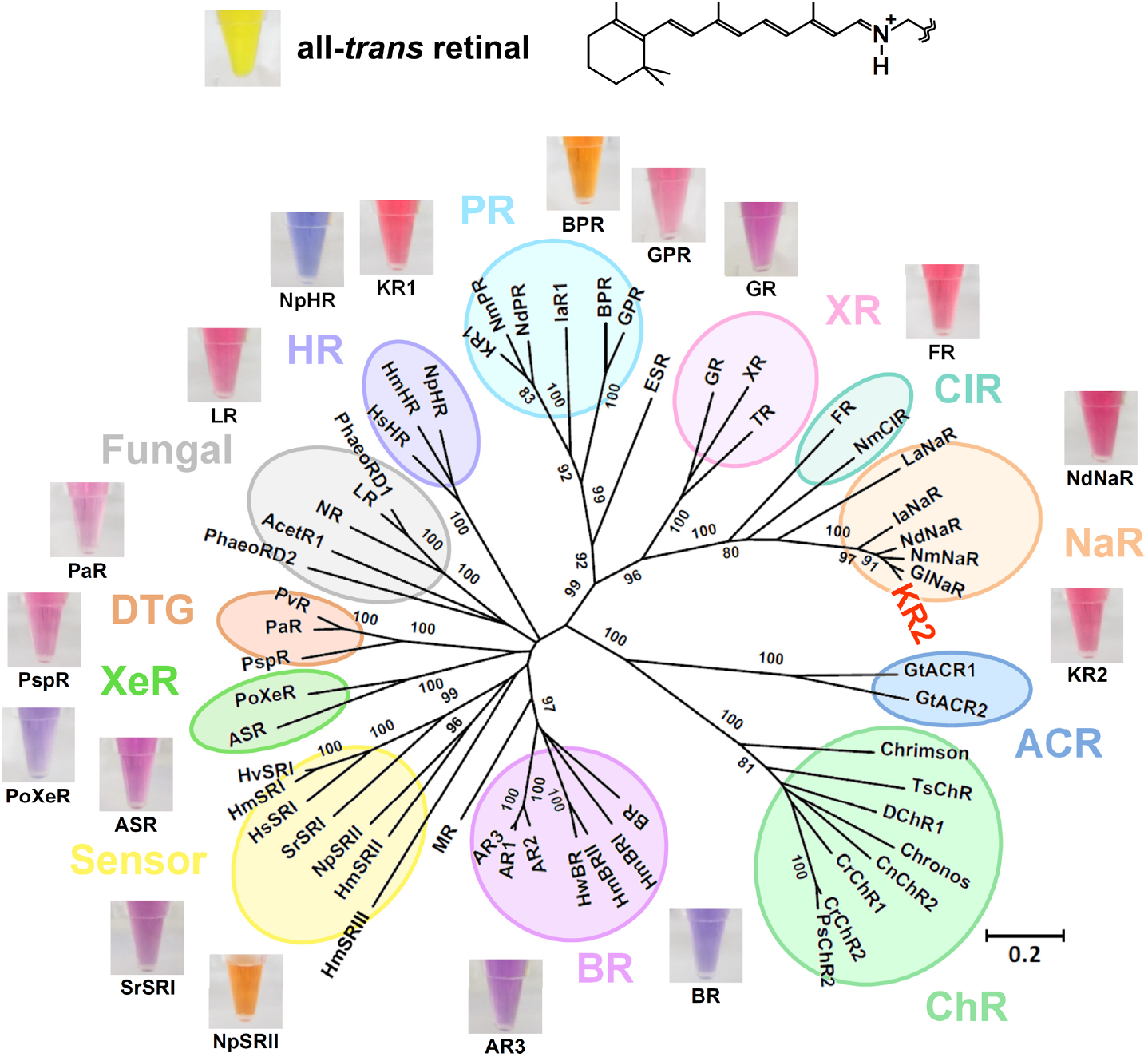
The chemical structure of all-*trans* retinal (upper) and phylogenetic tree of microbial rhodopsins (lower). The bootstrap values > 80% are shown for the corresponding branches. The photographs of the DMSO solution of all-*trans* retinal and detergent solubilized rhodopsins were aligned to show representative colours. The abbreviations of rhodopsin proteins are listed in Supplementary Information 1. In the present paper, we construct a machine-learning-based (ML-based) statistical model that describes the relationship between amino-acid sequences and absorption wavelengths of microbial rhodopsins based on past experimental data.

Increasing the variety of absorption wavelengths enables simultaneous optical control by different colours of light. Furthermore, the microbial rhodopsin having highly red-shifted absorption maximum is strongly demanded for optogenetic application, because of the lower phototoxicity and higher tissue-penetration length of longer-wavelength light^4^. As such, various rhodopsin genes have been screened in order to find additional colour-shifted proteins^4,6^. While many blue-absorbing rhodopsin at λ < 500 nm have been reported^7^ and even applied to optogenetics^4^, the longer absorption maxima are limited in < 600 nm. Thus, further artificial molecular modifications of protein were needed in order to achieve greater red-shifted absorption. Random and/or semi-empirical point mutations identify the types of amino-acid mutation that are effective for colour tuning^8,9^. Although numerous mutations causing bathochromic shift without disrupting protein function were identified in this way, the degree of shift is insufficient for application, and comprehensive screening is difficult because of the large number of possible mutations (> 20^200^). Although more rational molecular design is expected for quantum chemical calculation to estimate the absorption energy^10^, its high calculation cost makes application to wide-range screening difficult. An alternative technique for expanding the absorption range is the incorporation of natural or artificial retinal analogues^11^. For optogenetic application, however, a tissue-directed delivery method of these analogues must be developed.

In the present paper, we report the results of a data-driven approach for studying the light-absorption properties of microbial rhodopsin proteins by machine learning (ML). We constructed a database of 796 proteins consisting of microbial rhodopsin wildtypes and their variants, some of which were previously reported in the literature and others of which are newly reported herein (see Supplementary Table 1). Each entry of the database consists of the amino-acid sequence and absorption wavelength λ_max_ of a rhodopsin. We introduce an ML method for constructing a statistical model describing the relationship between amino-acid sequences and absorption wavelengths. The goal of the present paper is to demonstrate the effectiveness of ML-based data-driven approaches for functional protein studies. By constructing a database based on past experimental results and applying an ML method to the database, a statistical model describing the relationship between amino-acid sequences and molecular properties can be constructed. In the context of microbial rhodopsin studies, we illustrate the utility of such a statistical model by demonstrating that it can be effectively used for understanding the colour tuning rules and predicting the absorption wavelength (see Fig. 2).

**Figure 2:**
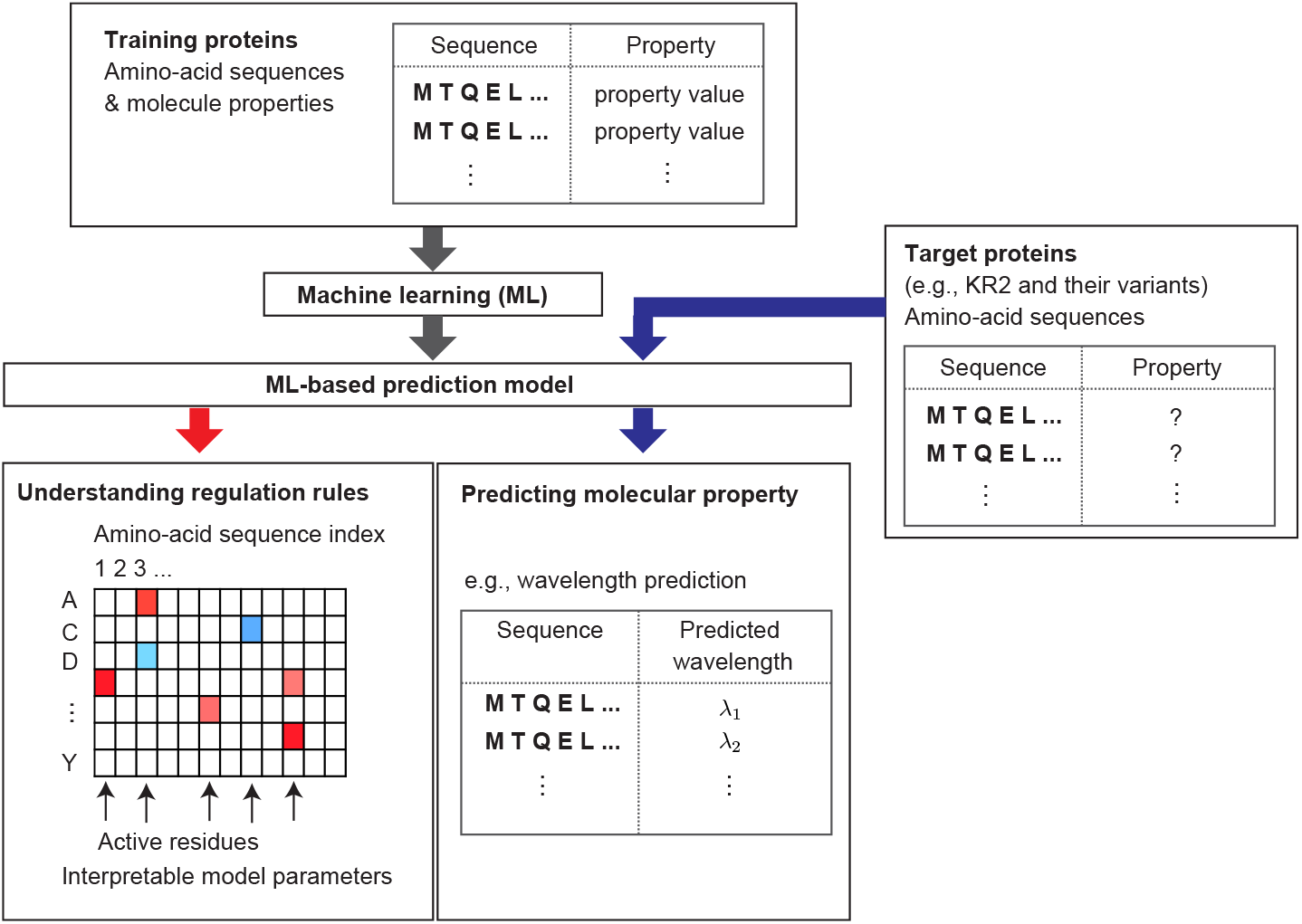
An overview of the machine-learning-based (ML-based) data-driven approach introduced in the present paper for functional protein studies. Using past experimental data, a *training protein set* containing pairs of amino-acid sequence and molecular properties is first constructed. Then, an ML method is applied to the training set, and an ML-based statistical model is constructed. The obtained ML model can be used in understanding the relationship between amino-acid sequences and molecular properties, such as the colour tuning rules in the case of microbial rhodopsins. The ML model can also be used to predict the molecular properties of new uninvestigated proteins. We refer to the set of new proteins as the *target protein set*. In the present paper, for the purpose of demonstration, we regard KR2 wildtype and its 118 variants as target proteins and other 677 rhodopsin proteins in the database as the training proteins.

We consider the following hypothetical scenario for the purpose of demonstration. The database is divided into two sets: a target protein set and a training protein set. The target set contains KR2 wild-type rhodopsin and its variants (which, in the present study, are assumed to be uninvestigated as of yet), whereas the training set contains the remaining proteins in the database. We constructed an ML model using only the proteins in the training set. The constructed model was then applied to the proteins in the target set for predicting the absorption wavelengths of KR2 and its variants. This scenario is interpreted as a hypothetical situation where a researcher is interested in predicting the absorption wavelengths of a new group of rhodopsin proteins based on previously reported data on other groups of rhodopsin proteins.

Among the various available ML methods, we used a *group-wise sparse learning* approach ^12,13,14^. The advantages of group-wise sparse learning approaches are not only predictability but also interpretability of the constructed models. As we report later herein, by using a group-wise sparse learning approach, the absorption wavelengths of KR2 and its variants could be predicted from their amino-acid sequences with an average error of ±7.8 nm. The residues affecting the absorption wavelength were also identified, and their strength for colour shift and the effect of mutation were quantitatively investigated. Through this analysis, the positions of BR Glu161 and Ala126, the effects for colour shift of which were not reported in previous studies, were newly shown to significantly affect the absorption wavelengths. Furthermore, the model constructed by a group-wise sparsity learning approach enables the identification of *active residues*, i.e., residues for which the choice of the amino-acid species has a great influence on the absorption wavelength. Although we herein focus on the prediction of absorption wavelengths of rhodopsin proteins, the same ML approach can be used to predict other molecular properties in other types of functional proteins.

## 2 Results

Microbial rhodopsin database In order to demonstrate the effectiveness of ML-based data-driven approaches for microbial rhodopsin studies, we constructed a database. The database is composed of amino-acid sequences and absorption wavelengths λ_max_s of 519 proteins previously reported in the literature and 277 proteins investigated by our group without previous report (see Supplementary Table 1). As reported in a previous study^15^, for data-driven approaches such as the present study, it is important to construct a database containing not only reported experimental results but also unreported results. We applied alignment algorithm ClustalW to these amino-acid sequences and obtained aligned sequences of 475 residues, among which we extracted the transmembrane region, resulting in 210 residues. For the purpose of demonstration, we divided the dataset into a *target protein set* and a *training protein set* (see Fig. 3).

**Figure 3:**
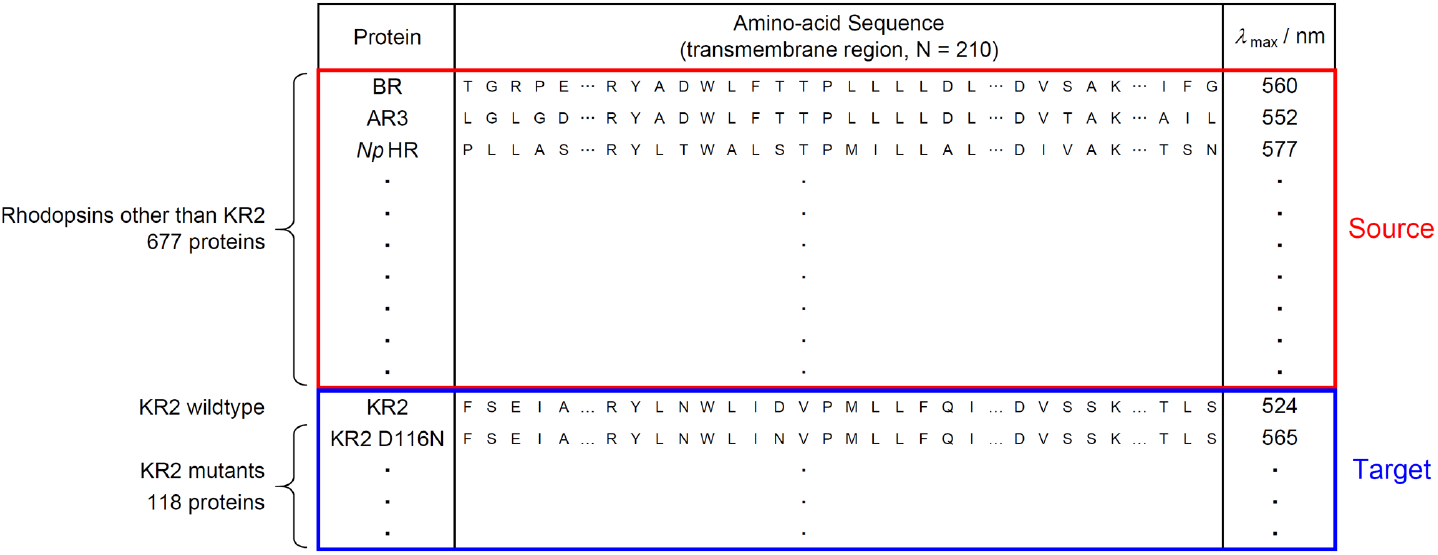
Structure of the database used in the present study. The database is composed of the sequences and λ_max_s of 519 previously reported proteins and 277 newly reported proteins. We used 677 rhodopsin proteins other than KR2 and their variants as the training proteins (red rectangle) and 119 proteins in KR2 group as the target proteins (blue rectangle), respectively.

The target set consists of 119 rhodopsin proteins in the KR2 group (KR2 wildtype and its 118 variants), whereas the training set consists of the remaining 677 rhodopsin proteins (see Figs. 1 and 3). We applied an ML method to the training set and constructed a statistical model describing the relationship between the amino-acid sequences and absorption wavelengths. The statistical model was then applied to the rhodopsin proteins in the target set in order to predict their absorption wavelengths. This scenario assumes a hypothetical situation in which a researcher is interested in investigating a new group of rhodopsin proteins based on previously reported data on other groups of rhodopsin proteins.

### Machine learning method

In order to handle amino-acid sequences in the ML framework, we introduced a binary representation, as depicted in Fig. 4(a). Let *M* = 20 be the number of different amino-acid species, and let *N* = 210 be the number of residues considered herein. Then, an amino-acid sequence is represented by *M* × *N* = 4, 200 binary variables, which we denote as ***x*** ∈ {0,1}^*MN*^. We consider a linear model for such *MN*-dimensional variables with an intercept parameter *β*_0_ and *MN* coefficient parameters *β_i,j_,i* = 1,…, *M,j* = 1,…, *N* (see Fig. 4(b)). These 1 + *MN* parameters are fitted based on the training set so that the output of the model *f* (***x***) can predict the absorption wavelength of the rhodopsin protein for which the amino-acid sequence is coded as *x*. Since this model has so many parameters, it is difficult to interpret the fitted model if we simply use conventional methods such as the least-squares method. We thus introduced the *group-wise sparsity mechanism* (See the Method section and the Supplemental information for details). Using this mechanism, the fitted coefficient parameters *β_i,j_* have *residue-wise sparsity*. Here, *M* = 20 coefficient parameters corresponding to the choice of an amino-acid species in each residue is considered as a group. After we fitted the model, in many groups, all of the *M* coefficient parameters become zero, indicating that the choice of an amino-acid species in these residues does not affect the colour tuning property. On the other hand, a small number of residues at which the coefficient parameters are NOT zero are called *active residues*, i.e., the choice of the amino-acid species in these residues is expected to play an important role in colour tuning. Figure 4(c) illustrates the fitted coefficient parameters using the group-wise-sparsity mechanism. If a parameter *β_i,j_* is positive/negative, then the *i*-th amino-acid species in the *j*-th residue has a red-shifting/blue-shifting effect on the light absorption properties of rhodopsin proteins.

**Figure 4:**
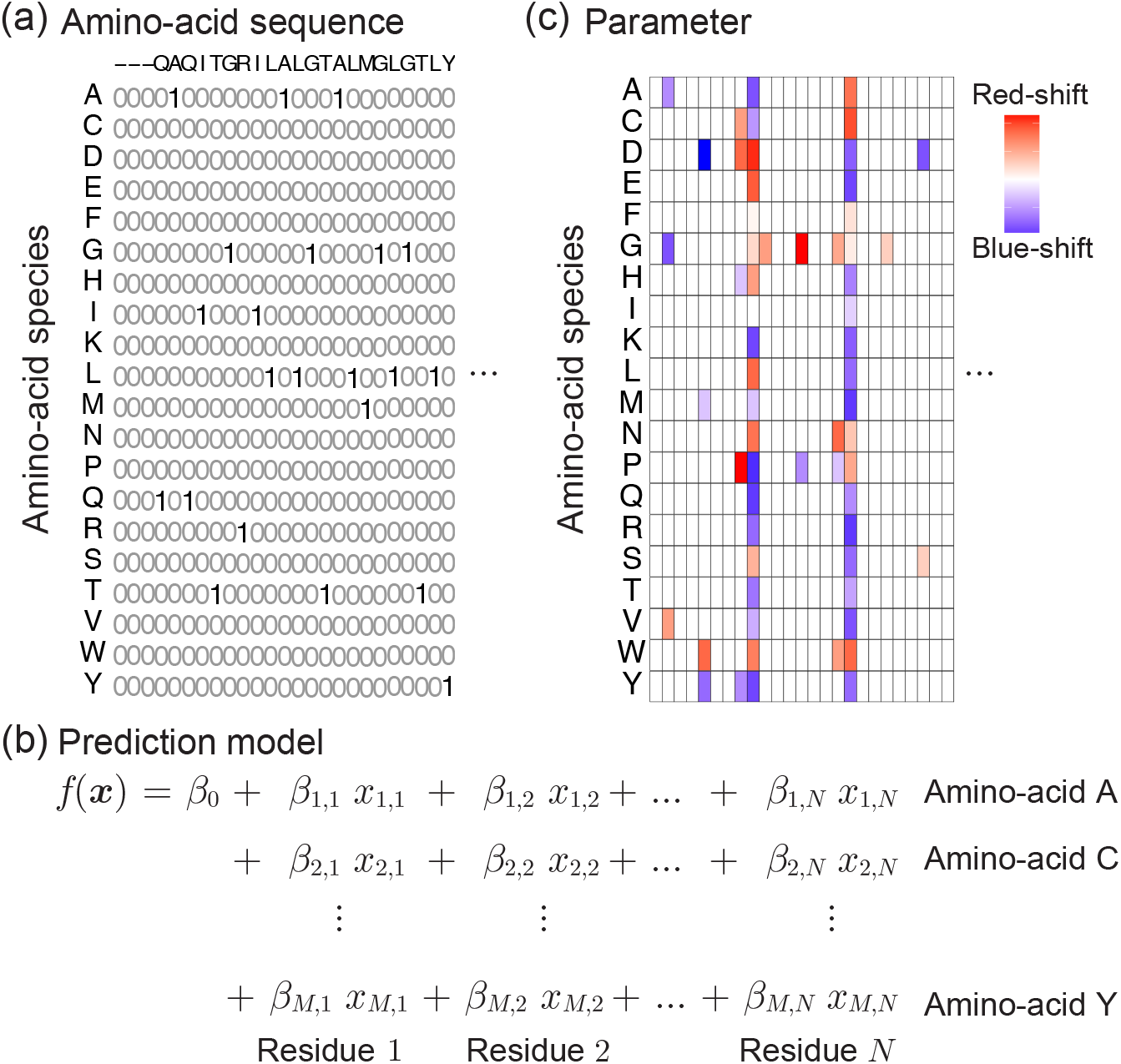
A schematic description of the ML method introduced in the present paper for functional protein studies. (a) Binary sequence representation of an amino-acid sequence. Let *M* = 20 be the number of amino-acid species, and let *N* be the number of residues considered in the present study. Then, the amino-acid sequence of a protein is represented by *M* × *N* binary variables, each of which represents the amino-acid species at each residue. (b) By writing the *MN* binary variables as *x_i,j_, i* = 1,…, *M,j* = 1,…, *N*, we consider an *MN*-dimensional linear model. The linear model has an intercept parameter *β*_0_ and *MN* coefficient parameters *β_i,j_, i* = 1,…, *M,j* = 1,…, *N*. (c) When the linear model is fitted, a group-wise sparsity constraint is introduced. Then, in many residues, all of the corresponding *M* coefficients would be fitted to zero, and only a small number of residues have nonzero coefficient parameters. The latter residues are called *active residues*. The choice of amino-acid species in these active residues is expected to play an important role in determining molecular properties such as absorption wavelength.

### Understanding colour tuning rules

By applying the above ML method to the training set containing pairs of the amino-acid sequence and absorption wavelength for 677 rhodopsin proteins, we fitted a linear model with 1 + *MN* = 4, 201 parameters. A complete list of the fitted parameters is presented in Supplementary Table 2. Figure 5 shows the fitted coefficient parameters at 20 active residues in decreasing order of 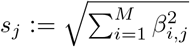, *j* = 1,…, *N*, where the score *s_j_* quantifies the *activeness* of the *j*-th residue. Here, red and blue indicate that the corresponding parameters are positive and negative, respectively, whereas grey indicates that the parameters were zero. In other words, red and blue suggest that having the amino-acid species in the residue would have a red-shifting and a blue-shifting effect, respectively. The results in Supplementary Table 2 and Fig. 5 can be interpreted as a comprehensive statistical description of the colour tuning rules of rhodopsin proteins based on previously investigated experimental results for 677 rhodopsin proteins (Supplementary Figure 1 shows the same results obtained using all 796 rhodopsin proteins, including those in the KR2 group).

**Figure 5:**
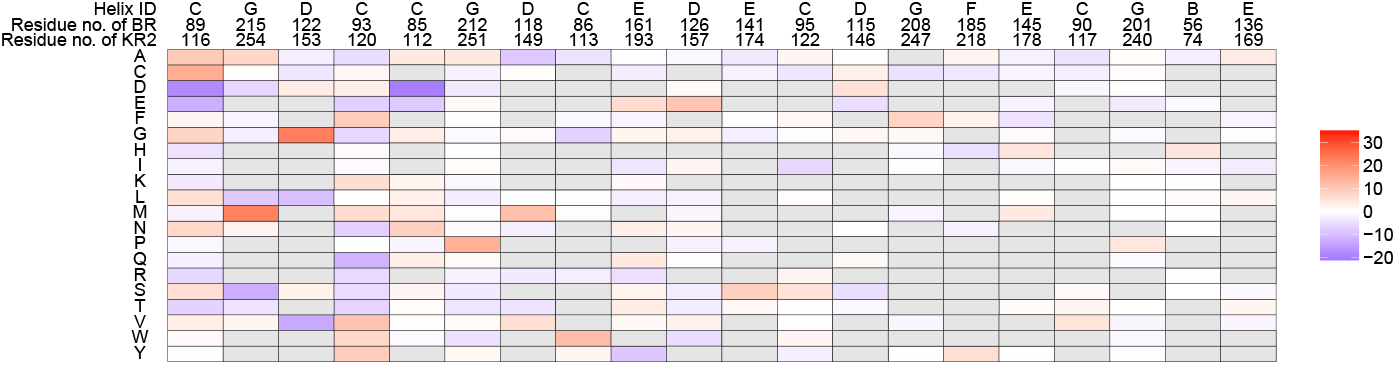
Coefficient parameters of the fitted statistical model. Coefficients for the top 20 active residues, where the activeness of each residue is defined as 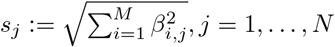. Here, red and blue indicate that the corresponding parameters are positive and negative, respectively, whereas grey indicates that the amino-acid species did not exist in the training data. The figure can be interpreted such that, if the value of a coefficient parameter *β_i,j_* is positive/negative (i.e., red/blue), then the existence of the *i*-th amino-acid species at the *j*-th residue has a red-shifting/blue-shifting effect.

### Predicting absorption wavelengths of KR2 rhodopsin and its variants

Using the statistical model fitted based on the training set (containing all of the rhodopsin proteins except for the KR2 group), the absorption wavelengths of the 136 rhodopsin proteins in the target set (containing KR2 group rhodopsin proteins) were predicted. Figures 6(a) and 6(b) show examples of predicted (green lines) and observed (blue lines) wavelengths for red-shifted KR2 mutants. For the KR2 NTQ/F72G mutant (Fig. 6(a)), the difference between the predicted (546.44 nm) and experimentally observed (543 nm) wavelengths is only 3.44 nm. In contrast, we observed a larger discrepancy (8.51 nm) for the predicted (556.49 nm) and experimentally observed (565 nm) wavelengths for KR2 D116N. This means that the precision of ML prediction differs for each type of mutation. Examples of blue-shifted mutants are shown in Figs. 6(c) (KR2 N112E) and 6(d) (KR2 DTD/D102N). The differences between the prediction and the observation were 7.34 and 19.92 nm for the former and latter, respectively. Figure 6(e) summarizes the prediction results for KR2 and all of its mutants, where the horizontal axis represents the *observed* absorption wavelengths measured in the experiments, whereas the vertical axis represents the *predicted* absorption wavelengths obtained by the ML model. The red points indicate the KR2 group rhodopsin proteins in the target set, whereas the black points indicate other rhodopsin proteins in the training set. Note that the prediction performance in the training set (black points) is slightly better than that in the target set (red points). This is because the former is used for fitting the ML model itself, whereas the latter is completely new to the fitted model. This phenomenon is known as *over-fitting* in the literature of machine learning. The absorption wavelengths of KR2 and its variants could be predicted from their amino-acid sequences with average errors of ±7.8 nm. The histogram in Fig. 6(b) shows the distribution of the prediction errors in the KR2 group rhodopsin proteins in the target set.

**Figure 6:**
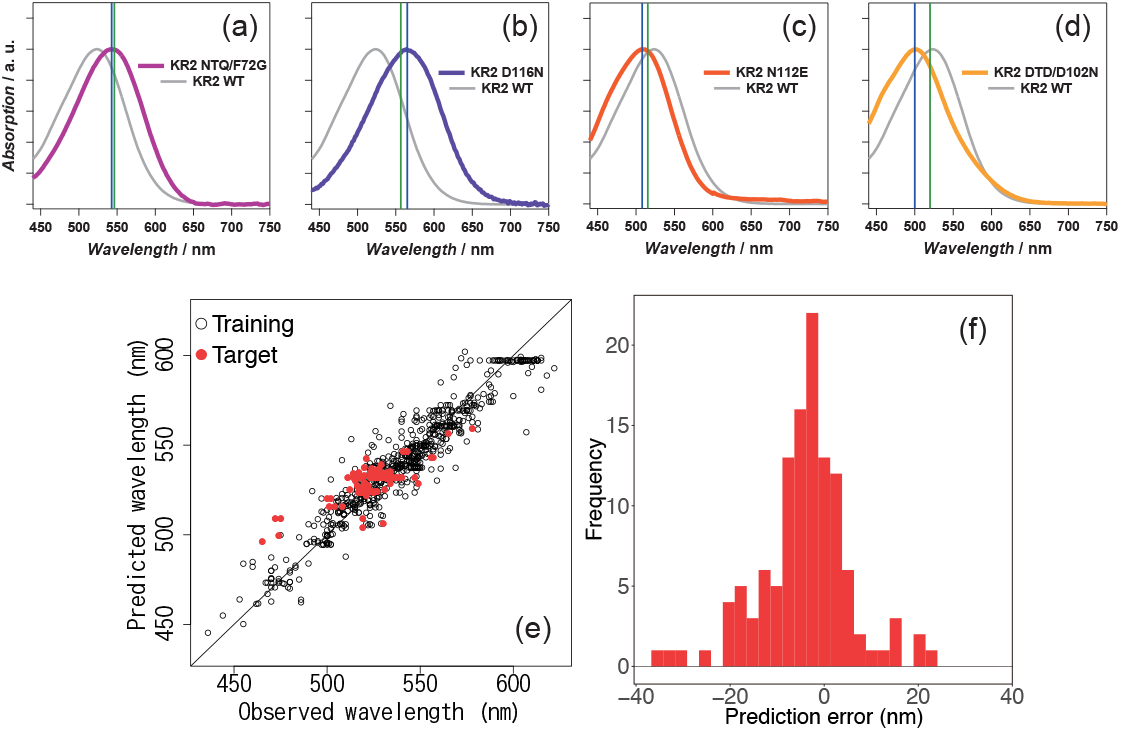
Absorption wavelength prediction results for KR2 wildtype and its 118 variants. (a)-(d) Absorption spectra of KR2 mutants ((a) KR2 NTQ/F72G, (b) D116N, (c) N112E, and (d) DTD/D102N) with their absorption maxima as predicted by ML analysis (green lines) and experimentally determined (blue lines). The spectrum of KR2 wildtype is indicated by the solid grey line. (e) The horizontal axis represents the experimentally observed absorption wavelengths, whereas the vertical axis represents the absorption wavelengths predicted by the ML model. The red points indicate the KR2 group rhodopsin proteins in the target set, whereas the black points indicate other rhodopsin proteins in the training set. (f) Histogram of the prediction errors for KR2 group proteins in the target set.

### Estimating the effect of point mutations

The effect of a point mutation on the absorption wavelength shift can be estimated based on the coefficient parameters *β_i,j_, i* = 1,…,*M, j* = 1,…, *N*. Let *x*^(KR2)^ ∈ {0,1}^*MN*^ be the binary vector representation of the KR2 wild-type sequence. The difference in the predicted absorption wavelengths between KR2 wildtype and a variant having amino-acid sequence ***x***^(Var)^ ∈ {0,1}^*MN*^ is written as

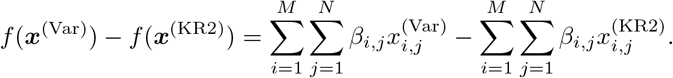

The colour-shifting effect of point mutation at the *j*-th residue is written as

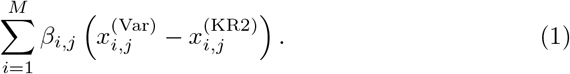

For example, if the *i*_1_-th amino-acid species in the KR2 wildtype is replaced by the *i*_2_-th amino-acid species, the colour-shifting effect of the point mutation is *β*_*i*2,*j*_ – *β*_*i*1,*j*_. Figure 7 shows a portion of the amino-acid sequences of KR2 wildtype and its variants along with their observed and predicted absorption wavelengths. In Fig. 7, red and blue indicate red-shifting and blue-shifting effects, respectively, in Eq. (1) estimated by the trained statistical model. Figure 7(a) suggests that point mutation at BR residue number 89 would have red-shifting effects. On the other hand, Fig. 7(b) suggests that point mutation at BR residues 85 and 122 would have blue-shifting effects. These results indicate that the estimated colour-shifting effects are consistent with the actual observed wavelength shifts caused by the mutation.

**Figure 7:**
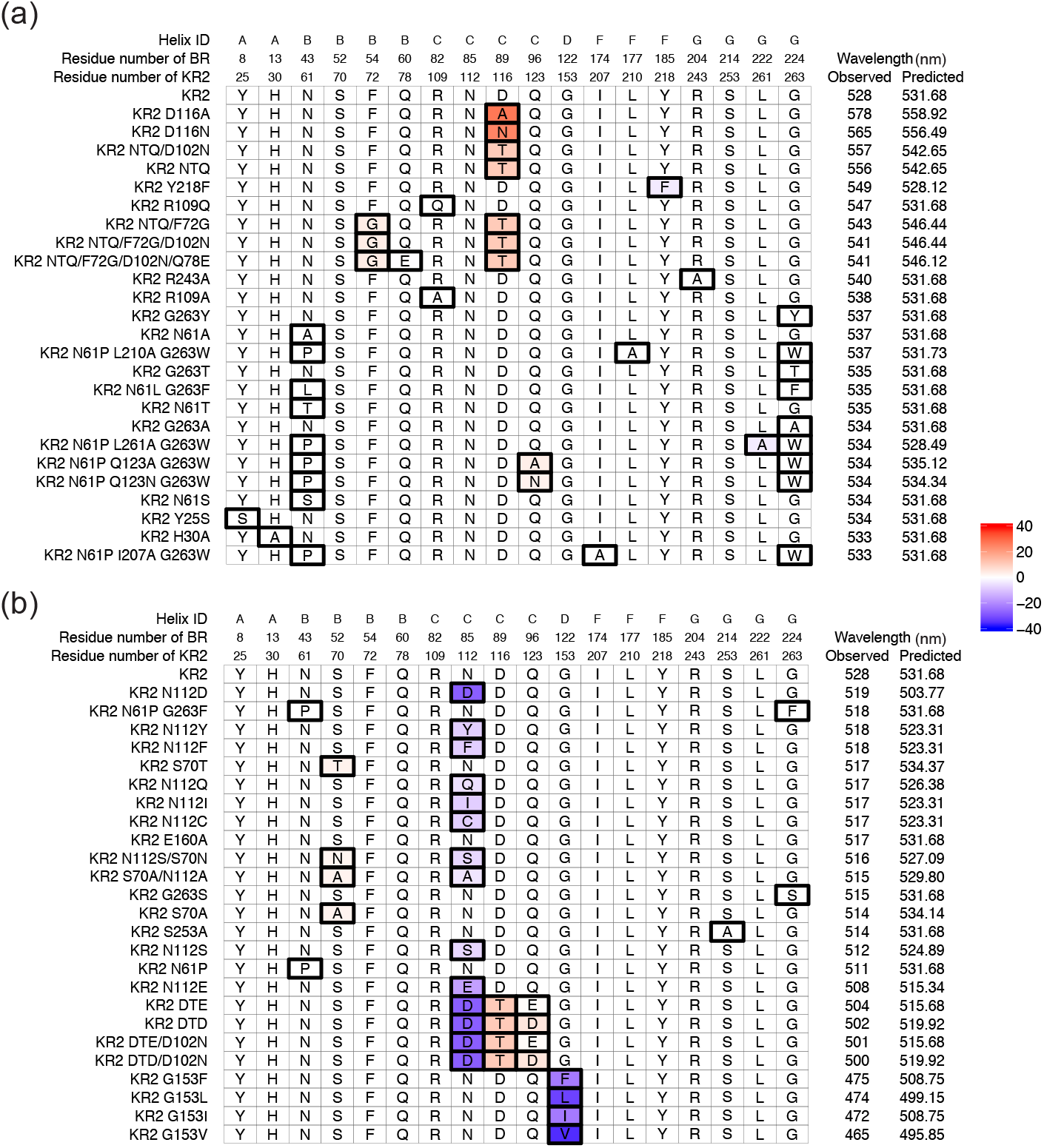
Lists of sequences for the KR2 wildtype and the variants with their observed and predicted absorption wavelengths. (a) KR2 and the 25 variants that have the longest observed wavelengths, and (b) KR2 and the 25 variants that have the shortest observed wavelengths. The residues shown here are replaced at least once among the 50 variants. Boxes with thick black lines indicate positions that have different amino-acid species from the KR2 wild-type. For these boxes, the colour indicates the wavelength change produced by the replacement of the *j*-th position, estimated by 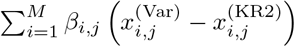, where 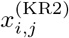 and 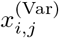 are the binary representation the KR2 wildtype and a variant, respectively.

## 3 Discussion

### Colour tuning rules in the estimated statistical models by ML

Ten residues showing the highest *β*-values were overlaid on the X-ray crystallo-graphic structure of BR (PDB code: 1BM1) (see Fig. 8). Eight of these residues are located around retinal within < 5 Å(BR Thr89, Ala215, Gly122, Leu93, Asp85, Asp212, Met118, and Trp86 in the order of degree of activeness). Thr89 showed the highest degree of activeness. This is a member of the DTD-motif, which represents the type of functional determining three residues in the third transmembrane helix (helix-C) for each ion-pump rhodopsin. The DTD-motif is typical for the outward H^+^ pump and is composed of Asp85, Thr89, and Asp96 for BR^16^. While this threonine is conserved among most microbial rhodopsins, it is replaced with an aspartate for sodium pump rhodopsin (NaR), which has the NDQ-motif rather than the DTD-motif ^17,16,18^. The position of BR Thr89 is close to RSB (the distance between BR Thr89C_*γ*_ and the nitrogen atom of RSB is 3.4 Å). The third and seventh active residues are BR Gly122 and Met118, respectively. These residues are highly conserved among various microbial rhodopsins. Their mutation causes the rotation of the C6-C7 bond of retinal and the shortening of the *π*-electron conjugation between the *β*-ionone ring and the polyene chain^19,20^. The largest coefficient parameters are obtained for glycine and methionine for the former and latter positions. This implies any type of mutation of these residues results in the blue-shift of λ_max_ and is consistent with previous experimental reports^18,19^.

**Figure 8:**
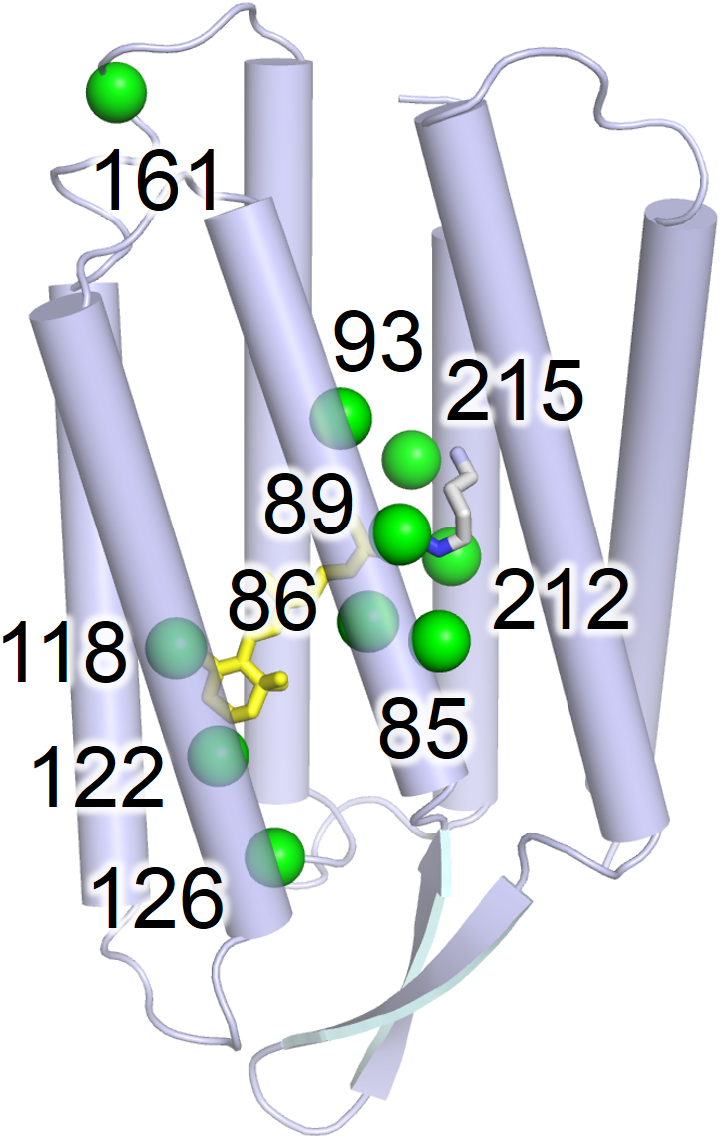
Top 10 active residues identified by the fitted statistical model. The positions of the active residues showing larger coefficient parameter values (green spheres) are mapped on the X-ray crystallographic structure of BR (blue, PDB code: 1BM1^27^) with their numbers in the case of BR.

The residues of BR Ala215 and Leu93 exhibit the second and fourth highest degrees of activeness. Both BR Ala215 and Leu93 are well known to have a role in colour-tuning switching for various rhodopsins in nature. Shimono et al. reported that, whereas green-to-orange absorbing archeal rhodopsins (BR, halorhodopsin and sensory rhodopsin I) conserve an alanine at the position of BR Ala215, blue-absorbing rhodopsins, such as *pharaonis* phoborhodopsin (*p*pR, which is also referred to as *pharaonis* sensory rhodopsin II) has a serine or threonine at this position^21^. The difference of coefficient parameter values is approximately 11.8, which is close to the reported λ_max_ shift of *p*pR T204A (8nm red-shift)^21^ and the BR homolog of *Haloquadratum walsbyi* (*Hw*BR) A223T (13-nm blue-shift)^22^. BR Leu93 corresponds to Leu120 of green-absorbing proteorhodopsin (GPR). This residue is replaced with a glutamine in blue-absorbing proteorhodopsin (BPR), and this type of colour regulation is known as “L/Q-switching”^23^. The lowest coefficient parameter (−11.2) was obtained for a glutamine. This suggests that glutamine is most effective to achieve blue-shift absorption and is considered to be optimized in natural evolution in the deep-ocean environment^23^. Ozaki et al. reported that mutations to valine or bulky residues (lysine, phenylalanine, tyrosine, and tryptophan) cause a large red-shift^24^ of λ_max_. Their larger coefficient parameters are consistent with previous experimental results (Fig. 5).

BR Asp85 and Asp212 are generally deprotonated and work as counterions to protonated RSB. The electrostatic interaction between their negative charges and the π-electron of retinal destabilizes the energy level of the electronically excited state. This results in the blue-shift of λ_max_^25^. Whereas the aspartate at the position of BR Asp85 has the second lowest coefficient value (−19.5) among all of the residues investigated herein, the value of the position of BR Asp212 is moderate (−3.2). This result suggests that the former has a much stronger effect on colour tuning, despite the symmetric location of these two residues relative to RSB. (The distances from Asp85 and Asp212 to the N atom of RSB are 3.4 and 3.5 A, respectively.)

The eighth largest coefficient parameter was the position of BR Trp86. This tryptophan is one of the most highly conserved residues among microbial rhodopsins. It forms a part of the binding pocket by direct contact with the extracellular side of the polyene chain of retinal1. This strong interaction with retinal is consistent with the high degree of activeness of this residue and the coefficient parameter of tryptophan is a large positive value (12.0). This suggests that this tryptophan has a role in shifting the absorption wavelength to be longer in many rhodopsins.

The positions of BR Glu161 and Ala126 are relatively far from retinal (having the 9-th and 10-th largest coefficient parameters). To our knowledge, there are no previous studies focused on the colour-tuning effects of these residues. For the position of BR Glu161, larger red- and blue shifts are expected for valine and tyrosine. In fact, sensory rhodopsin I (SRI), which is a positive phototactic sensor, has a valine at this position and exhibits relatively longer absorption maxima (e.g., the SRI of *Halobacterium salinarum* (*Hs*SRI): 587 nm; SRI of *Haloarcula vallismortis* (*Hv*SRI): 545 nm). In contrast, a tyrosine is conserved among various channelrhodopsins (ChRs), which generally have short absorption wavelengths (e.g., the ChR1 of *Chlamydomonas reinhardtii* (CrChR1): 453 nm; ChR1 of *Dunaliella salina* (*D*ChR1): 475 nm; ChR2 of *Proteomonas sulcata* (*Ps*ChR2): 444 nm). The results of ML analysis suggest the position of BR Glu161 is important for the colour tuning of these rhodopsins in nature. The position of BR Ala126 exhibited a large coefficient value for glutamic acid (10.5). Actually, *Gloeobacter* rhodopsin (GR), the outward H+ pump rhodopsin of cyanobacterium, *Gloeobacter violaceus* PCC 7421, has a glutamic acid at this position (GR Glu166), and the mutation of this residue exhibited a blue-shift of 1 to 22 nm (Supplementary Table 1). Thus, GR Glu166 works as an active residue for the colour tuning in GR.

These results imply the usefulness of ML analysis in identifying active residues located far from retinal, which are generally of less concern in experimental research on the colour tuning mechanism from a structural point of view. The effects on the absorption wavelength by the mutation of these residues have not yet been reported. However, we expect that they will be experimentally verified in the near future.

### Toward Experimental Design

The fitted linear model parameters *β_i,j_, i* = 1,…, *M, j* = 1,…, *N* can be also used as a guide for new functional protein design. For example, suppose that a researcher wants to construct a rhodopsin mutant, the absorption wavelength of which is as long as possible for opt-genetics application. Note that positive/negative coefficient parameter values indicate that the amino-acid species at the residue have a red-shifting/blue-shifting effect, respectively, on the light-absorption properties of rhodopsin proteins. Consider a residue *j* at which there exists *i*_1_ and *i*_2_ such that *β*_*i*1,*j*_ < *β*_*i*2,*j*_. If there exists a rhodopsin protein having the *i*_1_-th amino-acid species at the *j*-th residue, by replacing this species with the *i*_2_-th amino-acid species, the new protein is expected to have a longer wavelength than the original protein. This means that, the basic experimental design strategy for the above-mentioned researcher would be to replace the amino-acid species having a smaller coefficient parameter with that having a larger coefficient parameter. Although many other factors, such as protein stability and functionality, must be taken into account in new functional protein design, the above discussion suggests that the ML-based data-driven approach enables systematic design of experiments without relying on the intuition or heuristics of researchers.

## 4 Methods

### Construction of a dataset of amino-acid sequences and λ_max_s

For ML analysis, we constructed a database (Supplementary Table 1) composed of the amino-acid sequences and the previously and newly reported λ_max_s of microbial rhodopsins and their variants. Previously reported λ_max_s were collected from 102 reports (listed in Supplementary Information 2). Newly reported λ_max_s were experimentally determined in our group by the hydroxylamine bleaching method for *E. coli* membrane expressing rhodopsins^26^ or purified protein by Ni- or Co-NTA chromatography^17^, as described previously. The method used to determine each rhodopsin is also listed in Supplementary Table 1.

### Details of the ML method with group-wise sparsity regularization

Our data contains a larger number of variables (4, 200 binary variables) than the number of instances (677 rhodopsin proteins). In this case, classical least-squares methods may cause over-fitting of the training data, which results in poor prediction accuracy for the target data. *Sparse modeling* ^12,13^ is a standard approach to this problem setup so that only a small subset of coefficient parameters is automatically selected. In particular, we use a group-wise sparsity method^14^ to analyze the residue-wise effect on the absorption wavelength. Let *x_i,j_* ∈ {0,1} be a binary variable that indicates the existence of the *i*-th amino-acid species in the *j*-th residue, where *i* = 1,…, *M* and *j* = 1,…, *N*. Here, each *i* = 1,…, *M* of *x_i,j_* corresponds to one of *M* = 20 amino-acid species.

We consider predicting the absorption wavelength based on a linear model:

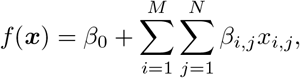

where *β*_0_ and *β_i,j_* for *i* = 1,…, *M* and *j* = 1,…, *N* are parameters. Suppose that we have *K* pairs of an amino-acid sequence and its absorption wavelength 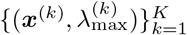, where 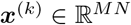 is the binary representation of the amino-acid sequence aligned as a vector, and 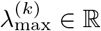 is the absorption wavelength of the *k*-th rhodopsin protein. The parameters are fitted by solving the following penalized least-squares problem:

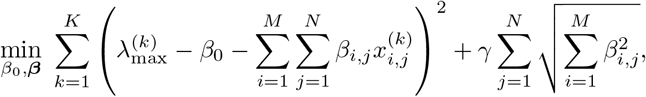

where *γ* > 0 is a tuning parameter. This formulation is called *group LASSO*^14^, in which the first term is the sum of the squared prediction errors, and the second term is the group-wise penalty for the parameters. For each residue *j* = 1,…, *N*, we define the *M* = 20 coefficient parameters (*β*_1,*j*_,…, *β_M,j_*) as a group. If the training set indicates that the choice of the amino-acid species at the *j*-th residue does not affect the colour tuning property, then the group-wise sparsity penalty forces all of the *M* = 20 parameters (*β*_1,*j*_,…, *β_M,j_*) to be exactly zero. We can easily identify a set of important residues for determining the absorption wavelength by this effect, called *group-wise sparsity*, because usually only a small subset of the residues have non-zero coefficient parameters. In our experiment, the parameter *γ* was objectively chosen by the cross-validation procedure within the training set.

### Code availability

Our program code of the group LASSO for wavelength prediction is available at http://…^1^

### Data availability

The database of the amino-acid sequences and their wavelengths is provided in Supplementary Table 1.

## Acknowledgements

We appreciate for the insightful discussion and data collection by Drs. R. Abe-Yoshizumi, M. Konno, Y. Kato, S. Ito, and Y. Inatsu. The present study was financially supported by grants from the Japanese Ministry of Education, Culture, Sports, Science and Technology to M.K. (16H06538 and 17H04694), K.I. (26708001, 26620005, and 17H03007), H.K. (25104009 and 15H02391), and I.T. (16H06538 and 17H00758), from JST PRESTO to M.K. (Grant Number JPMJPR15N2) and K.I. (Grant Numbers JPMJPR12A2 and JPMJPR15P2), from JST CREST to I.T. (Grant Numbers JPMJCR1302 and JPMJCR1502), from the RIKEN Center for Advanced Intelligence Project to I.T., and by the JST support program for starting up innovation-hub on materials research by information integration initiative to M.K. and I.T.

## Author contributions

M.K. analyzed the data by machine learning. K.I. constructed the database and interpreted the results. H.K. and I.T. designed the entire research study.

## Footnotes

1 The site will be public after acceptance. The code is attached to our submission.

